# Short-term Electrical Stimulation Impacts Cardiac Cell Structure and Function

**DOI:** 10.1101/2024.05.19.594880

**Authors:** Kristen Allen, Abigail Bandl, Natalie Pachter, Tracy Hookway

## Abstract

Induced pluripotent stem cell derived cardiomyocytes (iPSC-CMs) are used to model cardiac development and disease. This requires a robust population of mature CMs and external stimuli to mimic the complex environment of the heart. In effort toward this maturation, previous groups have applied electrical stimulation (ES) to CMs with varying results depending on the stimulation duration, frequency, and pattern. As such, there is uncertainty surrounding the timeline on which stimulated iPSC-CMs begin to show early signs of maturation in comparison to their non-stimulated counterparts, leaving room for additional research into when hallmarks of maturity–such as hypertrophy and changes to sarcomere structure–develop in CMs. Here, we introduce a low-cost custom bioreactor capable of delivering tunable electrical stimulation to standard 2D cell monolayers. We show that, after exposure to short-term ES, stimulated CMs express early signs of maturation compared to the non-stimulated control. The changes to contractility and protein expression indicate that the cells undergo hypertrophy in response to short-term ES, but they do not develop transverse-tubules, which is a key component of a fully mature CM structure. Therefore, while early signs of maturation are present after a short stimulation regimen, additional cellular structures must develop to reach complete maturation. We have shown that our custom electrical stimulator can be easily integrated with standard *in vitro* cell culture platforms to obtain measurable changes to cells, exhibiting its potential for promoting crucial CM maturation for cardiac tissue engineering applications.

## Introduction

The characterization of cardiac physiology and disease requires models of human *in vivo* tissue. However, discoveries made in murine, bovine, or porcine models often do not translate to human subjects (Paci et al., 2021). Because of these limitations, many groups have shifted to using *in vitro* models to model cardiac physiology and disease. One key roadblock to *in vitro* cardiac research is the availability of human cells and tissues. Induced pluripotent stem cells (iPSCs) offer a robust, ethical cell source for *in vitro* cardiac models, but there are still several concerns regarding the reliability of these models and whether the results are translatable to *in vivo* pathologies (Karbassi et al., 2020; Stoppel et al., 2016). Research is ongoing on the enhancement of iPSC-derived *in vitro* models to resemble *in vivo* tissue.

Robust differentiation protocols have been developed to differentiate iPSCs to cardiomyocytes (CMs), the contractile muscle cells within the heart (Lian et al., 2012). Stem cell-derived CMs are fetal in phenotype, featuring disorganized sarcomeres, spontaneous beating, less robust calcium handling, and a less defined shape (Karbassi et al., 2020). Long-term culture of iPSC-CMs without additional stimulation does not result in consistent maturation within a timeline feasible for disease modeling *in vitro*. Mature CMs boast an organized sarcomere structure, synchronized beating with neighboring cells, and a rod-like cell structure (Bedada et al., 2016). These characteristics mirror adult cells *in vivo* and are ideal for studying CM gene expression, structure, and ion handling.

Other groups have employed various methods to induce maturity of fetal-like iPSC-derived CMs, including electrical (Gabetti et al., 2023; Ye & Black, 2014), mechanical (Karbassi & Murry, 2022; Morin et al., 2014; Ye & Black, 2014), and chemical stimuli (Abecasis et al., 2019; Yang et al., 2019). Electrical stimulation is popular due to the role of electrical signals in the cardiac system. To begin a beat, an electrical signal is sent from the SA node to the AV node between the right atrium and ventricle (Figure 1a) (Marieb & Hoehn, 2010). Upon receipt, the signal travels from the AV node through the bundle branches and subendocardial conducting network to coordinate the beat of the ventricular myocardium (Arthur et al., 1971; James, 2001). These signals establish different cardiac rhythms, including normal sinus rhythm and diseased states such as bradycardia and ventricular tachycardia (Figure 1b) (Kalra et al., 2018). Electrophysiology plays a crucial role in CM behavior *in vivo*, motivating the exploration of electrical stimulation on *in vitro* cardiac tissue models.

**Figure 1:**
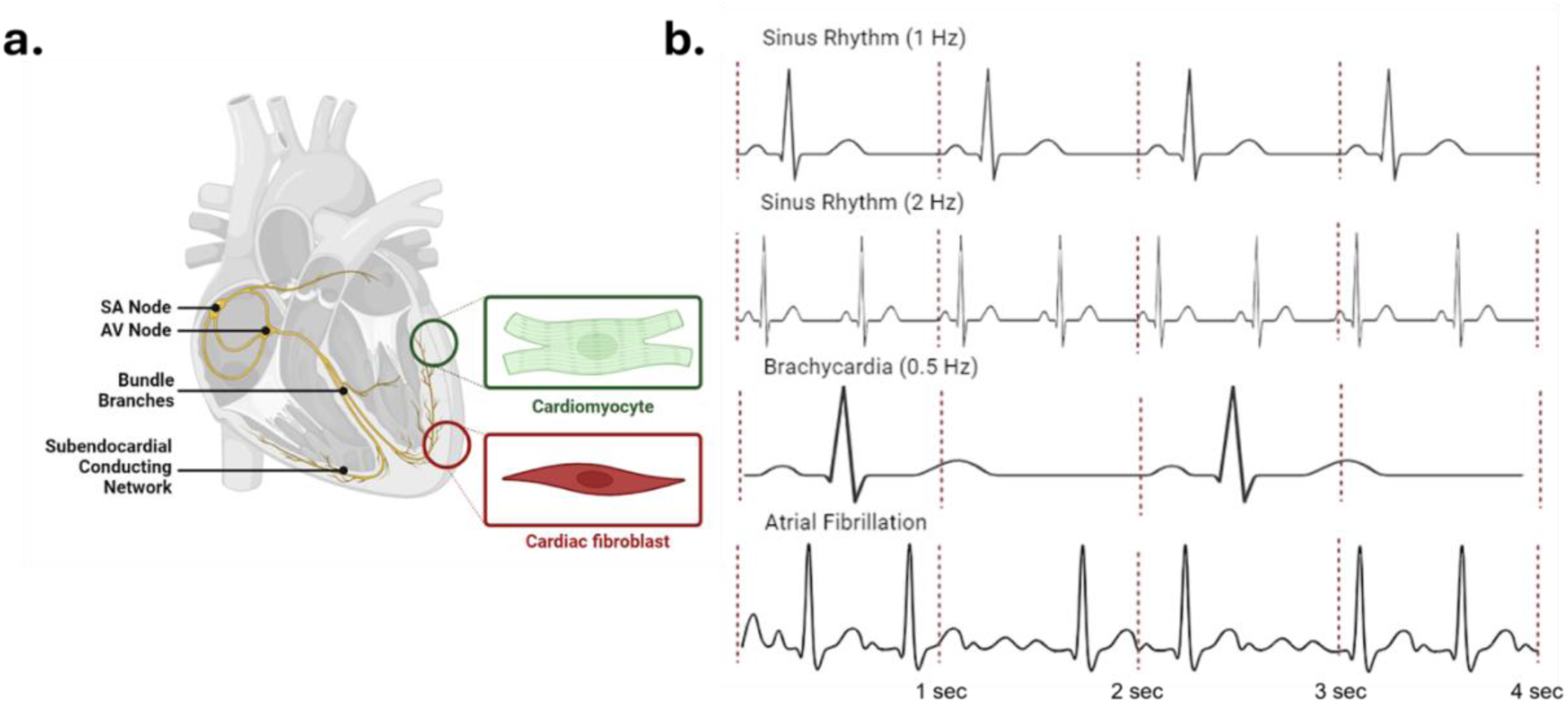
Importance of electrical stimulation in the heart. **a)** Electrophysiological schematic of heart anatomy *in vivo*, **b)** physiologically relevant cardiac rhythms with the red dotted lines indicating 1 second of time.

Electrical stimulation has been applied with varying results depending on the stimulation duration, frequency, and pattern (Eng et al., 2016; Nunes et al., 2013; Ronaldson-Bouchard et al., 2018; Sesena-Rubfiaro et al., 2023). There is uncertainty surrounding the timeline on which stimulated iPSC-derived CMs achieve maturation in comparison to their non-stimulated counterparts, with room for additional research into when hallmarks of early maturation–such as hypertrophy, changes to sarcomere structure, and transverse-tubules–develop in CMs. Additionally, many electrical stimulation systems are costly and difficult to use. Here, we have developed a low-cost electrical stimulation bioreactor that fits on a standard 6 well plate and provides tunable rhythmic and arrhythmic electrical stimulation. The device can be easily integrated into existing cell cultures and can provide 2 different stimulation speeds and rhythms within the same plate. Using this bioreactor, we observed changes in both morphology and function in CMs and cardiac fibroblasts (cFB) upon variable frequency electrical stimulation.

## Methods

### Differentiation of iPSC-CMs

Two lines of WTC11 iPSCs, one genetically encoded with a GCaMP-6F calcium reporter (Gladstone Institutes) and one with a cardiac troponin I (TNNI) reporter (Allen Institutes) were differentiated to CMs. Cells were plated at a density of 46k cells/cm^2^ on Matrigel-coated tissue culture plates and differentiated to CMs by modulating the Wnt pathway, yielding committed and spontaneously beating CMs by Day 10 (GCaMP, Lian et al., 2012) or Day 14 (TNNI, Gerbin et al., 2021; Grancharova et al., 2021). Upon differentiation, CMs were replated for electrical stimulation. Briefly, CMs were washed twice with PBS, leaving the second wash on for 18 minutes at room temperature to aid dissociation. Cells were then dissociated with Trypsin (Corning) for 10 minutes at 37°C, then counted and replated onto Matrigel-coated 6 well polystyrene plates at 300-475,000 cells/cm^2^ (GCaMP) or 270,000 cells/cm^2^ (TNNI). The replated cells formed cohesive monolayers and resumed spontaneous beating within 4 days of the replate. CMs were fed with RPMI/B27+ media supplemented with 1% penicillin-streptomycin (PenStrep) every 3 days.

### Cardiac Fibroblast Culture

Primary adult human cardiac fibroblasts (cFB) (Cell Applications) were thawed and seeded on tissue culture-treated 6-well polystyrene plates at 42,000 cells/cm^2^. The cells were fed with commercial cardiac fibroblast media (Cell Applications) or in-house cardiac fibroblast media (DMEM/F12, 10% fetal bovine serum, 1% L-glutamine, 1% nonessential amino acids, 1% PenStrep) every 2 days until reaching confluence.

### Custom Electrical Stimulation Bioreactor Design

The device consists of a 6 well plate lid 3D printed (Formlabs) in Formlabs BioMed clear resin (Figure 2b). Protruding from the inner part of the lid are 12 rod-shaped carbon electrodes, designed so 2 electrodes sit in each well of a 6 well plate at 25 mm apart, similar to the design featured in Mobini et al., 2016. The lid is attached to an electrical box featuring an Arduino UNO microprocessor and control circuit (Figure 2a). Work is done through the cell culture media when the Arduino microprocessor switches the state of the transistor, allowing for current to flow between the electrodes. To impart a voltage density of 1.15 V/cm throughout the cell culture media, 2.8 V of DC power is supplied to the circuit through the power supply (Au et al., 2009).

**Figure 2:**
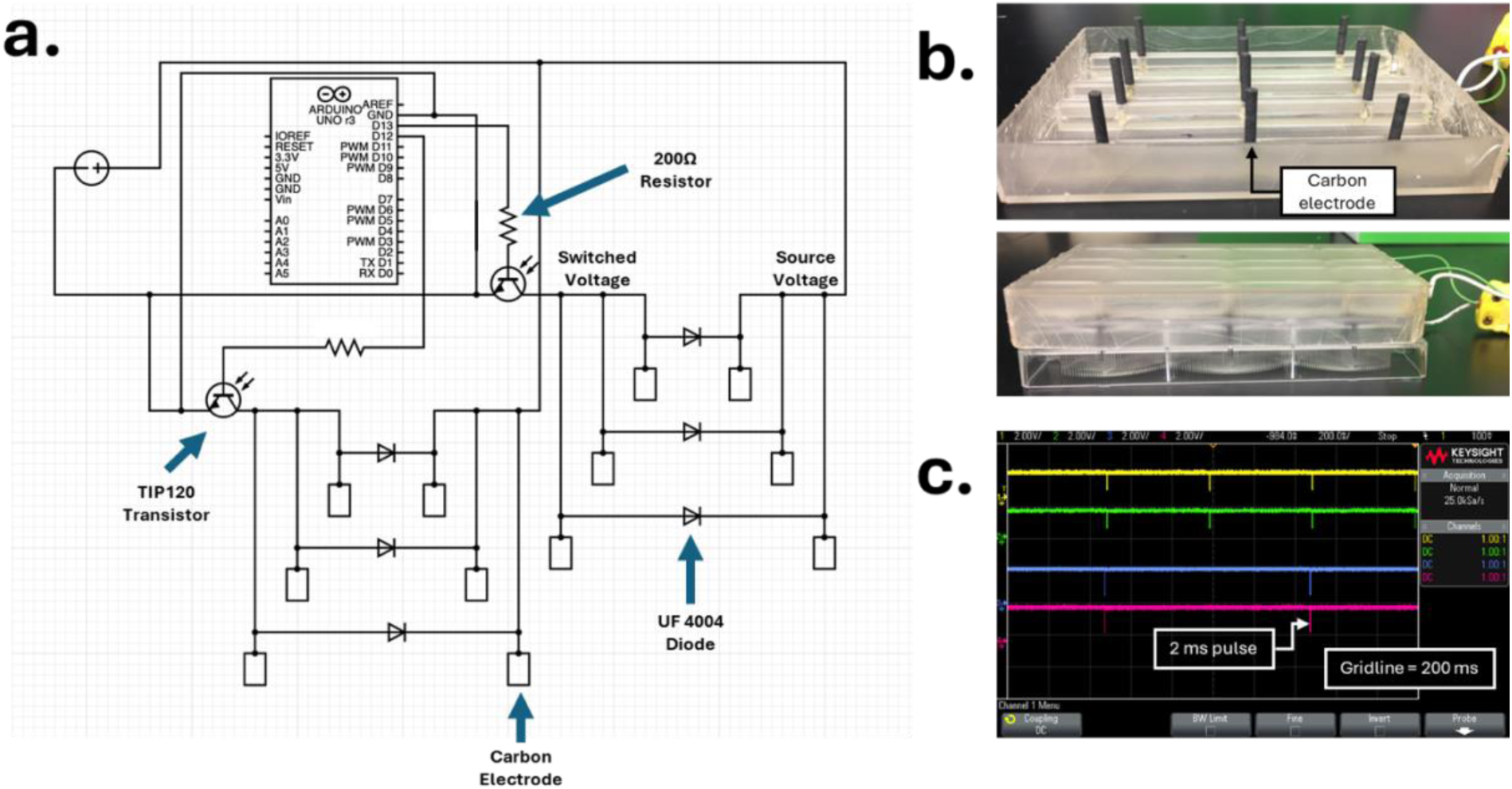
Design and verification of the custom electrical stimulation device. **a)** Circuit schematic, **b)** custom lid to standard 6 well cell culture plate, **c)** oscilloscope reading of “switched voltage” side of UF4004 diode and corresponding electrode (yellow = 2 Hz diode, green = 2 Hz electrode, blue = 1 Hz diode, red = 1 Hz electrode).

The device is designed to run any electrical stimulation pattern desired by the user via editing of the Arduino code. In this study, the device was set to provide 2 ms square monophasic pulses because this mirrors the stimulation provided to the myocardium *in vivo*, and the 2 ms stimulation is long enough to generate an excited response from the CMs (Marieb & Hoehn, 2010; Nuccitelli, 1992; Tandon et al., 2009, 2011). The device imparted 0 Hz (unstimulated control), 1 Hz, and 2 Hz stimulation patterns to monolayer cultures of CMs or cFBs (Figure 2c). Due to resin deformation in autoclave, the device was sterilized in a 70% ethanol bath for 20 minutes, washed with cell culture grade water, then exposed to UV light for 30 minutes.

### Electrical Stimulation

CMs were allowed to recover for 4 days after replating, then exposed to 0 (unstimulated control), 1, and 2 Hz of electrical stimulation for 3 (GCaMP) or 4 (TNNI) days (Eng et al., 2016). cFB, once confluent, were exposed to the same electrical stimulation parameters for 4 days.

### Contractility and Beat Kinetic Assessments

On Days 0, 1, and 3 of stimulation, stimulation was paused for 1 hour for live imaging. The cells were placed on a heated stage (Ibidi Silver Line) for the acquisition of high-quality phase videos (Nikon Eclipse T*i*2 with equipped Andor Zyla camera). Five regions of interest (ROIs), each measuring 100x100 µm, were captured for each of the 3 wells per condition. The high-quality videos were converted from ND2 to AVI format using a custom ImageJ macro. The AVIs were then processed by MUSCLEMOTION, an ImageJ Plugin used to analyze beat kinetics (Sala et al., 2018; Schindelin et al., 2012; Schneider et al., 2012). MUSCLEMOTION contraction profiles were manually checked to ensure all contractile peaks were captured by the software. An ROI was considered compromised if an artifact was present in the video or if the ROI was not beating during capture. A well was included in analysis if it retained at least 2 non-compromised ROIs. The outputs were averaged across a well, resulting in 3 independent samples per condition and parameter within a single experiment, with most samples derived from an average of 4-5 ROIs.

### Immunostaining

After 1 day of recovery from stimulation, cells were fixed with 4% paraformaldehyde and washed 3 times with PBS. TNNI CMs were stained with Hoechst (1:4500) for 20 minutes. Images were acquired on a Nikon Ti2 Eclipse microscope with Andor Zyla camera. The green fluorescent cardiac troponin I (cTnI) expression of the CMs was measured through mean intensity using ImageJ (Schindelin et al., 2012; Schneider et al., 2012) and further quantified by thresholding, binarizing, and measuring the percent coverage of the green fluorescence (cTnI).

Cardiac FB were stained with phalloidin (1:50) and Hoechst (1:4500). Actin filament alignment based on phalloidin staining was assessed with DirectionalityJ, a built-in package for ImageJ (Schindelin et al., 2012; Schneider et al., 2012). DirectionalityJ is programmed using the local gradients orientation algorithm, which utilizes the Laplacian of the Guassian filter of an image to identify boundaries (Liu et al., 1991; Sensini et al., 2018). These boundaries are then used to identify linear regions (i.e. actin filaments) and assess their directionality (Liu et al., 1991). Only directionality measurements with a “goodness of fit” (GOF) value greater than 0.5 were retained for further quantitative analysis. Data below this threshold was excluded because a low GOF value indicates that the orientation measurements taken in the image were highly variable. This variability in directionality could not be reliably represented by a single directionality value, which would have skewed results of quantitative analysis.

### Statistical Analysis

All data was assessed for normality and homogeneity using the Shapiro-Wilks Test and Levene’s Test, respectively. For data containing one variable, all of the groups displayed normality and homogeneity, and one-way ANOVA was utilized to assess statistical significance. For data containing 2 variables, the data was found to be homogeneous, but not normally distributed. Despite the lack of normality, 2-Way ANOVA was still run to elucidate how increasing time in culture and stimulation group impacted the cells concurrently. This is considered a gold-standard in the field, and 2-Way ANOVA is an extremely conservative statistical test for data that is not normally distributed, more so than any non-parametric alternative (Feys, 2016).

## Results

### Hardware and Software Verification of Electrical Stimulation Device

The hardware and software of the electrical stimulation bioreactor were verified by oscilloscope readings taken on the “switched voltage” side of the diode and matching electrode (Figure 2c). This test verified that work was being done across the electrodes at a pulse width of 2 ms through the voltage drop measured on the “switched voltage” electrode. It further verified that stimulation was occurring at the correct frequency of 1 Hz and 2 Hz, and the correct voltage density of 1.15 V/cm was being imparted through the liquid, as that was the amplitude of the voltage drop seen in Figure 2c. Finally, the equivalence of the diode and electrode measurements verified the UF4004 diode was a viable test point for what was occurring across the electrodes. Validation of this diode means users can test if the correct stimulation protocol is being run in the device during cell culture studies without compromising the sterility of the experiment.

### Compatibility of Cardiomyocyte Culture is Dependent on Voltage and Seeding Density

Monolayers of CMs were exposed to 2.92 V/cm (7.3 V from the DC power supply) or 1.15 V/cm (2.8 V from power supply) (Au et al., 2009; Ronaldson-Bouchard et al., 2018). Upon exposure to 2.92 V/cm, the CM 2D monolayers experienced extensive detachment in response to two days of stimulation (Figure 3a). This indicated that this high voltage density was not compatible with the monolayers of CMs seeded on polystyrene. When the voltage density was decreased to 1.15 V/cm, CMs maintained cohesive, beating monolayers (Figure 3a), indicating that the lower voltage was compatible with 2D culture on polystyrene (Au et al., 2009).

**Figure 3:**
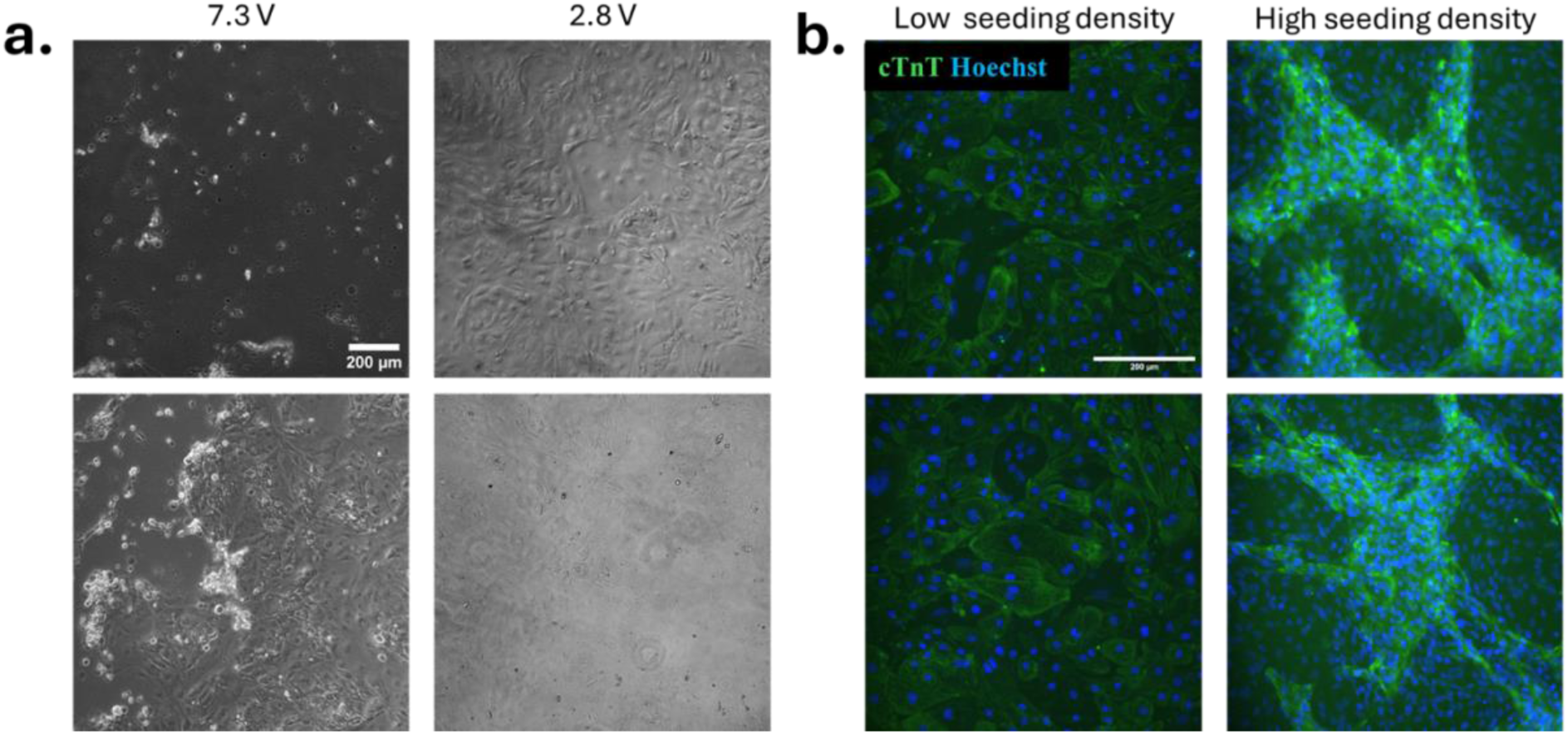
Biological verification of custom electrical stimulation device. **a)** Comparison of cell attachment in response to electrical stimulation at two imparted voltages, **b)** comparison of cell attachment in CM monolayer cultures at two different seeding densities in response to stimulation at 2.8 V. Scale bars = 200μm.

Upon confirmation of a compatible voltage density for CMs, further investigation was conducted into the impact of cell density on response to electrical stimulation. The need for such investigation was elucidated by the cell detachment observed directly under the carbon electrode when the monolayers were formed from a low seeding density of 67,700 cells/cm^2^. The monolayers seeded at this low density were compromised in the areas under the electrode, and the CMs did not arrange into connected, uniformly beating regions. However, at higher seeding densities of 270,000-300,000 cells/cm^2^, cohesive monolayers developed as pictured in Figure 3b, with dense cohesive beating patterns within the syncytium. Additionally, the monolayers formed from the higher seeding density were not compromised in the regions directly below the carbon electrodes. Therefore, it was determined that the higher seeding density was required for cell testing in this electrical stimulation bioreactor, and subsequent studies with CM monolayers were conducted at 270,000-300,000 cells/cm^2^.

### Cardiac fibroblast culture is compatible with electrical stimulation device

We next aimed to elucidate how electrical stimulation impacts cardiac fibroblasts (cFBs), since these cells are also abundant in the native heart and are important for propagation of electrical signal throughout the heart. Given the effect of cell density observed with CMs, cFBs were seeded and grown to confluence prior to initiating electrical stimulation. Upon exposure to 1 Hz of stimulation for 4 days at 1.15 V/cm, cFBs remained attached in culture and exhibited robust actin expression as seen through phalloidin staining (Figure 4a,b). There was a slight decrease in actin expression in the 2Hz condition, but cells maintained a complete monolayer in all conditions, indicating the compatibility of the stimulation device with additional cell types, including cFBs.

**Figure 4:**
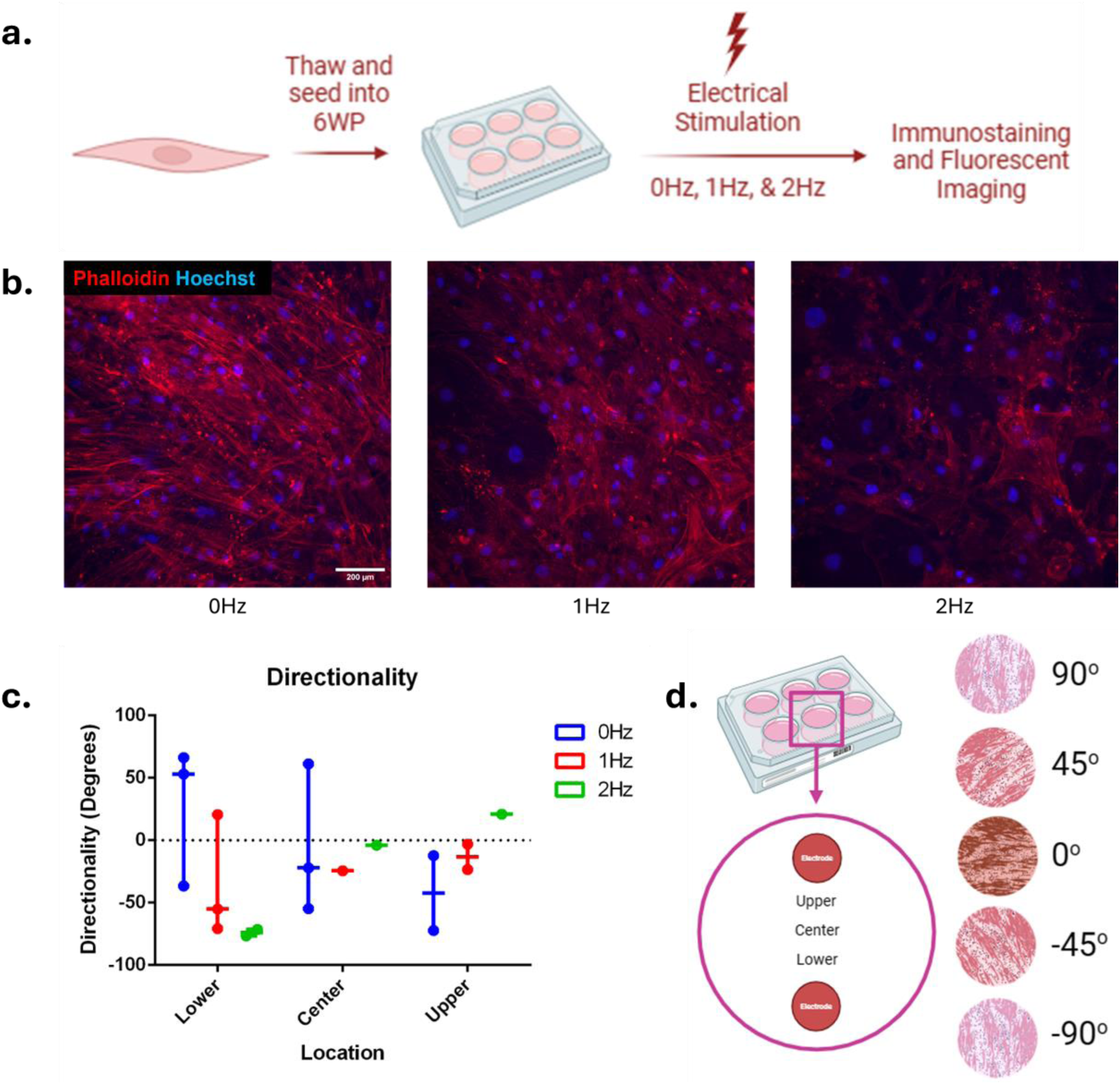
Custom electrical stimulation device is compatible with cardiac fibroblasts. **a)** Electrical stimulation experimental workflow on cFBs, **b)** fluorescent images of phalloidin and Hoechst staining of cFBs exposed to stimulation (scale bar = 200 µm), **c)** directional orientation of cFBs sorted by location in the well, **d)** schematic of the well locations and orientations.

Analysis of the directionality of the actin expression was used to determine if the cFBs reorganized their structural arrangement in response to electrical stimulation at the same frequency and voltage densities as CMs. After 4 days of 0, 1, and 2 Hz stimulation, no discernable trend in cFB alignment was seen at the center of the well, however, the greatest rearrangement in directionality was observed in the upper and lower locations, which were closest to the electrodes (Figure 4c,d). Although not significant, the stimulated groups, particularly the 2 Hz group, exhibited greater alignment near the electrodes. This can be seen by the 2 Hz group angled at -80° by the lower electrode, whereas the fibroblasts naturally arranged in the positive direction in the lower region of the 0 Hz condition. Likewise, the 2 Hz group oriented in the positive direction near the upper electrode when the 0 Hz group naturally arranged on the plate in the negative direction. These observations suggest that the directionality of the fibroblasts near the electrodes diverges from the directionality of the non-stimulated group. This response indicates that cFBs are compatible with the device, and additional cardiac cell types could be used in future studies to investigate the joint impact of electrical stimulation and co-culture on CM maturity.

### CMs exhibit structural changes under electrical stimulation

After four days of electrical stimulation, a robust population of CMs remained attached to the cell culture vessel (Figure 5b). The comparison of the mean intensity of fluorescently-tagged cTnI across the stimulation groups elucidates a significant difference between the 0 Hz and 2 Hz groups, indicating a stronger expression of sarcomeric proteins in the stimulated CMs (Figure 5c). In contrast, the percent coverage of the fluorescent signal significantly decreased in the stimulated groups compared to the 0 Hz control (Figure 5d), suggesting fewer cells. Taken together, these measures indicate that although there are fewer CMs contained in the stimulated cultures, the expression of the sarcomeric organization is higher in the CMs exposed to 2 Hz stimulation.

**Figure 5:**
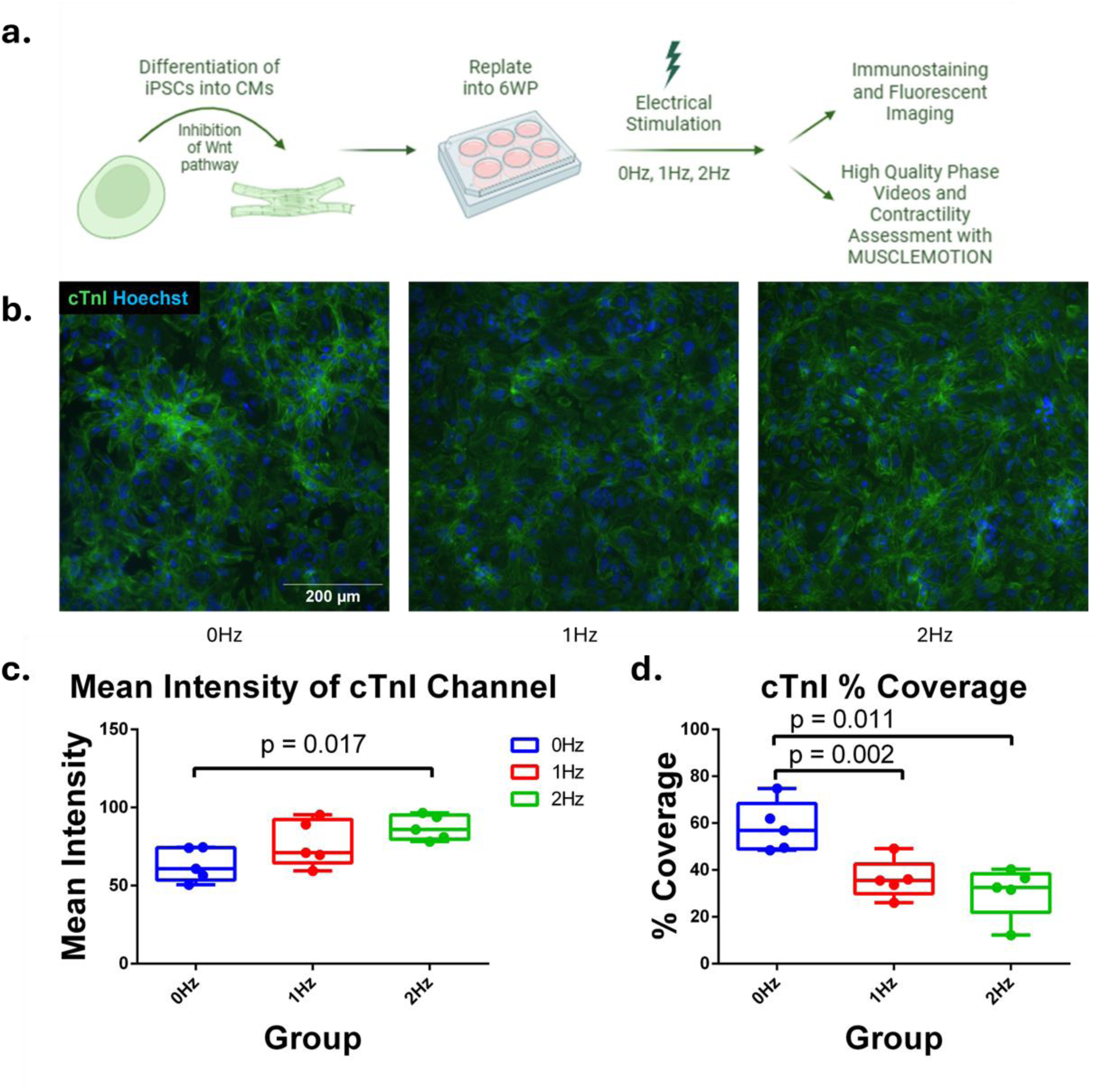
Robust monolayers of cardiomyocytes undergo structural changes after exposure to electrical stimulation. **a)** Electrical stimulation experimental workflow on CMs, **b)** fluorescent images of cTnI expression and Hoechst staining of CM (scale bar = 200 µm), expression of cTnI measured by **c)** mean intensity in a fluorescent image and d) percent coverage.

### iPSC-derived CMs express changes in contractility after exposure to short-term ES

To determine any functional consequences of short term electrical stimulation on iPSC-CMs, contractility assessment was conducted on four populations with pooled results shown in Figure 6. After exposure to 2 Hz of stimulation for 3 days, the beat frequency, contraction amplitude, and beat shape of CMs changed. This is highlighted by the significant increases seen in the peak-to-peak time, contraction duration, time-to-peak, and relaxation time (Figure 6a, d-f) and the significant decrease in beats per minute (Figure 6b) of the 2Hz group on day 3 compared to day 0. Although not significant on day 3, similar trends in the data were observed with the 1 Hz stimulation group as well. These changes indicate that increased stimulation frequency led to physiological changes in CMs by reducing beat frequency and prolonging contractions, nascent signs of maturation.

**Figure 6:**
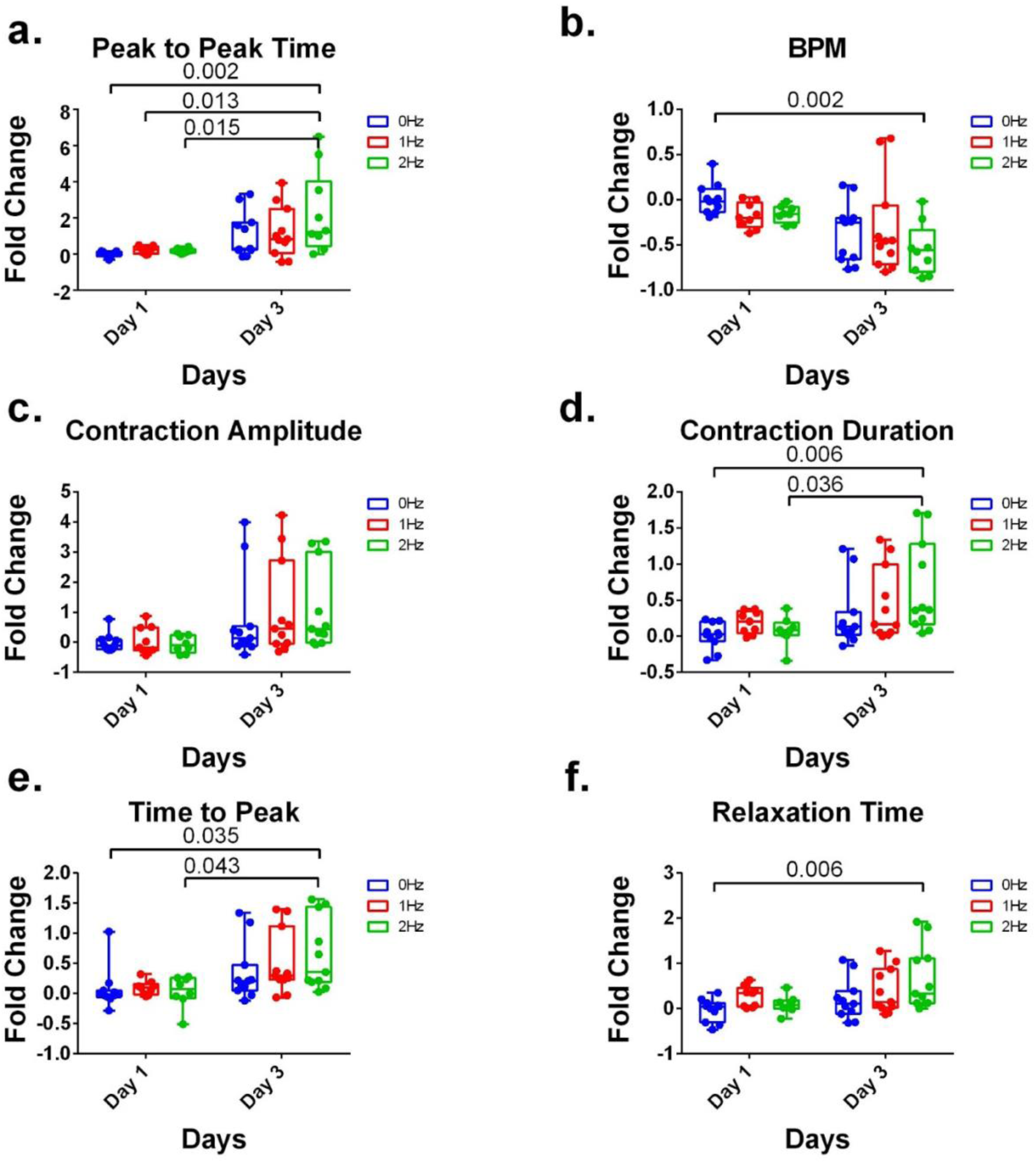
Significant changes to contractile parameters in response to time in culture and stimulation frequency. Fold change in **a)** peak to peak time, **b)** beat frequency (beats per minute) **c)** contraction amplitude, **d)** contraction duration, **e)** time to peak, and **f)** relaxation time when compared to Day 0 values from 2D monolayer of iPSC-derived cardiomyocytes under electronic stimulation at 0 Hz, 1 Hz, and 2 Hz for 3-4 days.

**Figure 7:**
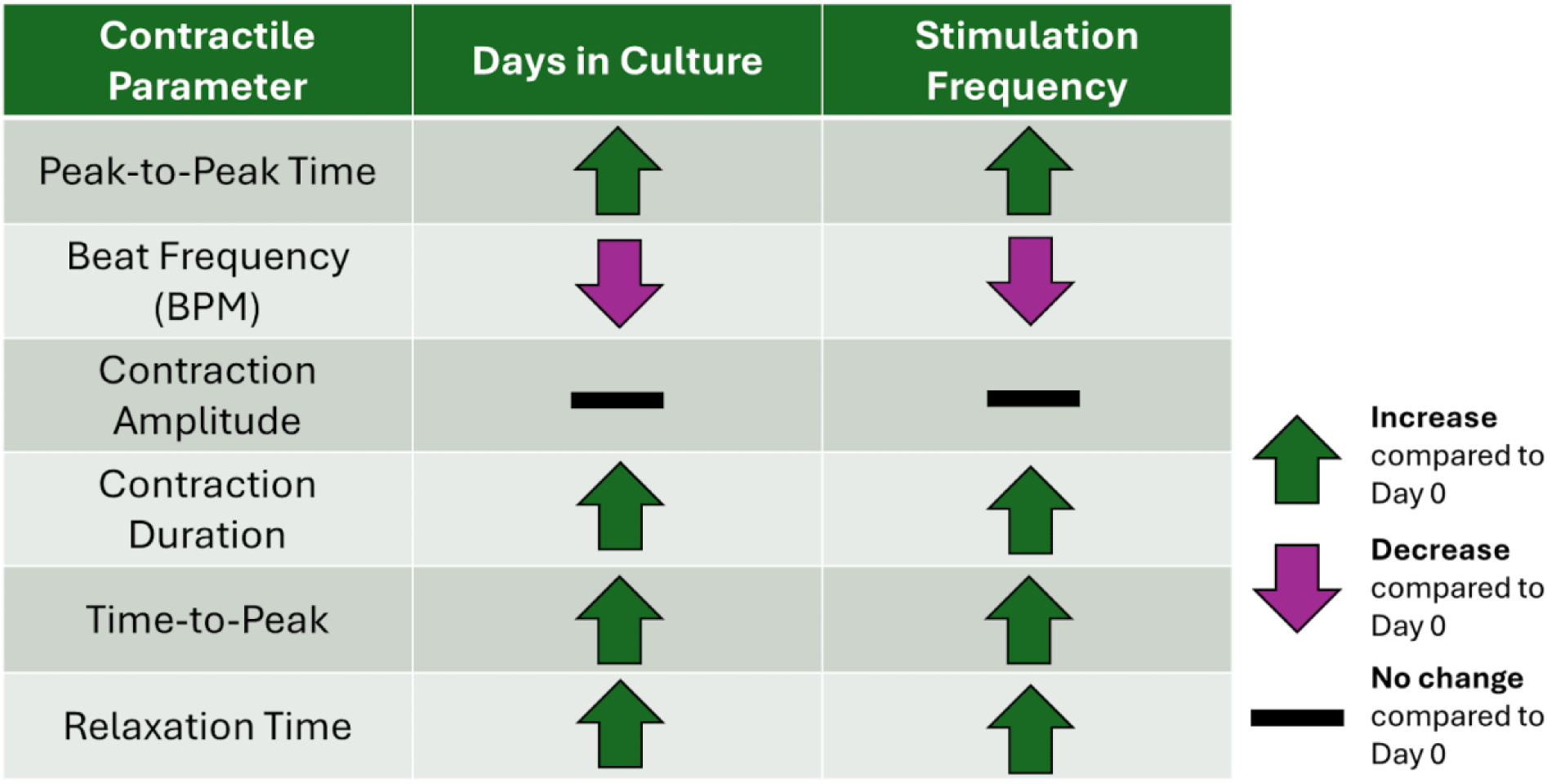
Summary of changes to contractility within iPSC-derived CMs. Trends observed in response to increasing stimulation frequency and days in culture when compared to day 0 (before stimulation).

## Conclusion

In this study, we designed a custom electrical stimulation bioreactor that was used to subject different 2D cardiac cell populations to varied stimulation frequencies. This low-cost stimulation device offers advantages not available in other devices seen in the literature. Utilizing a standard 6-well plate format allows for seeding either monocultures as described in this study and coculture experiments, limiting the confounding variables that can be present in custom chambers necessary in other devices. The device is also easily sterilizable and can also be expanded to other well sizes, with adjustments to the imparted voltage based on the distance between the electrodes. Finally, the device presented here is inexpensive compared to commercially available electrical stimulation devices, making it far more accessible to laboratories of any scale with access to a 3D printer.

There are, however, limitations to the device presented. It is likely that the voltage density is not consistent throughout the well, meaning that the cells receive varied intensity of stimulation based on location (Gabetti et al., 2023). This is likely one reason why increased fibroblast rearrangement was observed closer to the electrodes compared to the center of the well (Figure 4c). However, despite this limitation, the voltage density variation can be accounted for in the analysis phase by averaging across the well, ensuring that this device still accomplishes the ultimate goal of applying controlled stimulation to cardiac cell cultures as a method of promoting maturation. Therefore, the benefits of the device, including ease of sterilization, customizability, and cost-effectivity, outweigh the limitation of variation in the exposed CMs. The data presented indicates the device is effective at inducing a degree of maturation, and further work will allow for more advanced maturation of CMs.

In this study, the bioreactor was used to assess changes in cellular structure and function in response to short-term electrical stimulation, modeling normal sinus rhythm. The CMs exposed to electrical stimulation experienced a decrease in beat frequency and an increase in the duration of the beat. These beat kinetic changes commonly occur in iPSC-derived CMs that display pathological hypertrophy; however, healthy physiological hypertrophy also occurs as fetal-like cardiomyocytes mature into adult CMs *in vitro*, yielding the beat kinetic changes seen in the data presented (Frey & Olson, 2003; Li et al., 2022; Nunes et al., 2013). Further, physiological hypertrophy also explains why the number of CMs decreased in the stimulated groups, as indicated by the significant decreases in the percent coverage of cTnI+ cells, while the mean intensity expression of cTnI increased in the 2 Hz stimulation group (Figure 5c-d). While some cells detached in response to the stimulus, the cells that remained attached appear to have experienced hypertrophy, increasing the expression of cTnI as the sarcomeres expanded in a larger cell volume.

Likewise, as hypertrophy occurred in the stimulated cultures, the cell size increased, increasing the distance between interior sarcomeres and the sarcoplasmic reticulum, which is the primary storage location for intracellular calcium (Guo & Pu, 2020). In fully developed cardiac tissue, this distance is closed by the existence of transverse-tubules (t-tubules), which bring Ca^2+^ ions directly to the interior sarcomeres of the CMs (Guo & Pu, 2020). However, fetal-like iPSC-derived CMs do not contain organized networks of t-tubules, meaning the CMs lack the t-tubule infrastructure to deliver Ca^2+^ from the sarcoplasmic reticulum directly to the interior sarcomeres (Ormrod & Ehler, 2023; Satin et al., 2008). Instead, residual calcium is delivered to the interior sarcomeres through the “calcium-wave” propagating via calcium induced calcium release (CICR) (Satin et al., 2008). Hypertrophy, without t-tubule formation, causes delayed arrival of intracellular calcium ions to interior sarcomeres because the calcium signal must propagate further, decreasing the contraction frequency (Satin et al., 2008). Fully developed iPSC-derived cardiac tissue would be characterized by hypertrophy, but also by the development of t-tubule structures, which would increase the beat frequency despite the longer distance intracellular calcium must travel. The CM monolayers in these experiments exhibited the increased peak-to-peak time and decreased frequency characteristic of hypertrophy and incongruent with t-tubule formation. These CMs were in the transient state between fetal and adult CMs and therefore exhibited signs of both mature and immature CMs.

The partial maturation of CMs in response to short-term electrical stimulation captures a novel time point that is often overlooked in the literature; however, it also highlights the question of when full maturation occurs in response to stimulation. Extending the stimulation protocol in stages would also allow for identification of the time point at which cells experience t-tubule formation, which would be captured by an increase in beat frequency and through transmission electron microscopy (Guo & Pu, 2020; Miki et al., 2021; Sesena-Rubfiaro et al., 2023). Further, the formation of t-tubules may also be supported through the introduction of cFBs in co-culture with CMs (Ormrod & Ehler, 2023). Cardiac fibroblasts remained attached to the culture plate using the custom stimulator lid (Figure 4b) and remodeled in response to electrical stimulation (Figure 4c). This means further investigation is required into whether iPSC-derived CMs would mature faster when exposed to electrical stimulation in a CM-cFB co-culture. The next step in enhancing *in vitro* modeling is to develop a robust pipeline for the availability of robust, mature, and pure populations of CMs; electrical stimulation, in combination with co-culture and CM purification protocols, will likely play a key role in accomplishing this.

## Acknowledgements

We would like to thank the Watson Fabrication Lab at Binghamton University for their help in printing the custom lid. All schematics are produced in Biorender. Funding for this study was provided by the National Science Foundation (CAREER EBMS 2237898) and National Institutes of Health (1R15HL140745).

## Author Contribution Statement

K.A. and A.B. designed the stimulation device; K.A., N.P, A.B, T.A.H conceptualized and designed experiments; K.A. and N.P performed experiments, data collection, and analysis; K.A., N.P., and T.A.H were responsible for data interpretation and manuscript preparation; T.A.H was responsible for obtaining funding and project supervision.

## Ethics Statement

The authors confirm that the study and reporting contained here adheres to ethical guidelines as listed by the journal.

